# Benefits of siderophore release lie in mediating diffusion limitation at low iron solubility

**DOI:** 10.1101/093948

**Authors:** Gabriel E. Leventhal, Martin Ackermann, Konstanze T. Schiessl

## Abstract

Siderophores are chelators released by many bacteria to take up iron. In contrast to iron receptors located at the cell surface, released siderophores are at risk of being lost to environmental sinks. Here, we asked the question whether the release itself is essential for the function of siderophores, which could explain why such a risky strategy is widespread. We developed a reaction-diffusion model to determine the impact of siderophore release on overcoming iron limitation caused by poor solubility in aerobic, pH-neutral environments. We found that secretion of siderophores can efficiently accelerate iron uptake at low solubility, since secreted siderophores solubilize slowly diffusing large iron aggregates to small, quickly diffusing iron-siderophore complexes. At high iron solubility, however, when the iron-siderophore complex is no longer considerably smaller than the iron source itself, siderophore secretion can also slow down iron uptake. In addition, we found that cells can synergistically share their siderophores, depending on their distance and the level of iron aggregation. Overall, our study helps understand why siderophore secretion is so widespread: Even though a large fraction of secreted siderophores is lost, the solubilization of iron through secreted siderophores can efficiently increase iron uptake, especially if siderophores are produced cooperatively by several cells.

## Introduction

Iron is important for bacterial cell growth and reproduction, but iron availability is limited in many environments. One common cause of iron limitation is low concentrations, such as in the oceans (Boyd & Ellwood, 2010). But iron availability even at higher concentrations can also be poor, due to the physicochemical properties of iron. More specifically, low iron solubility has been described as a widespread cause for iron limitation in aerobic, pH-neutral environments (Braun & Killmann, 1999; Kraemer, 2004).

One of the various strategies bacteria employ to acquire iron is siderophore secretion, a widespread and well-studied mechanism (Hider & Kong, 2010). Siderophores are chelators that bacteria release into the environment to bind iron. The resulting iron-siderophore complexes are then again taken up by the bacteria.

An important consequence of secretion is that a cell might not recapture and thus benefit from siderophores it produced, due to random diffusion of the siderophore molecules. In dilute, well-mixed environments, the probability of recapturing a siderophore once it is secreted is low, and a solitary bacterium thus has to produce a large number of siderophores in order to achieve sufficient uptake of iron (Völker & Wolf-Gladrow, 1999). Also, released siderophores can be taken up by strains that do not contribute to siderophore production if these express the cognate receptor (De Vos *et al.,* 2001; West & Buckling, 2003). This can lead to a public goods dilemma, where nonproducing genotypes can displace bacteria that produce siderophores (Velicer, 2003).

Bacteria can also acquire iron with alternative mechanisms that avoid the disadvantages of secretion. For example, *Pseudomonas mendocina* can acquire iron upon direct physical contact with an iron-containing mineral by surface-associated reductases (Kuhn *et al.,* 2013). Many bacteria also use outer membrane receptors for the uptake of iron bound to exogenous chelators like heme or transferrin (Andrews *et al.,* 2003). Similarly, ferric citrate can be taken up via transporters or porins (Marshall *et al.,* 2009). It has also been suggested that siderophores can stay attached to the cell, e.g. in some marine bacteria (Martinez *et al.,* 2003) or at conditions of low cell density (Scholz & Greenberg, 2015).

Despite the existence of such alternative mechanisms that avoid the risk of siderophore loss, siderophore secretion is widespread in bacteria (Sandy & Butler, 2009), and has been described as key for iron uptake in environments with low iron availability (Miethke & Marahiel, 2007). This suggests that the *release* of siderophores might be directly beneficial for their function. However, it is less clear what possible benefits might be. In order to identify and quantify these benefits, we developed a mathematical model that enables us to compare the efficiency of iron uptake with and without the release of siderophores. We hypothesize that secretion of siderophores is especially important at low iron solubility, where diffusing siderophores can help overcome diffusion limitation caused by large, slowly moving iron aggregates that form due to poor solubility. Siderophores can solubilize these iron aggregates, generating quickly diffusing iron-siderophore chelates, potentially significantly speeding up iron uptake.

## Results

### Development of a model to describe varying iron solubility

As a first step to constructing a model for siderophore-iron interaction, we develop a description of the iron distribution at varying solubility. In aerobic, pH neutral environments, the predominant iron species, ferric iron, is poorly soluble. Low iron solubility results in the formation of iron aggregates (Kraemer, 2004), and one of the primary reported functions of siderophores is the solubilization of such iron aggregates (Vraspir & Butler, 2009). Iron aggregates are polymorphous, containing iron as well as hydroxide or other groups. Over time, and depending on external conditions like pH, the aggregates change in crystal structure and size (Cornell *et al.,* 1989; Schwertmann *et al.,* 1999). Here, we simplify this complexity by assuming that iron aggregates are spherical, and we account for varying solubility of iron by adjusting the number of iron ions, k, that comprise an aggregate. Soluble iron corresponds to the smallest iron aggregate, i.e. one iron ion surrounded by water molecules (*k* = 1). The number of iron ions *k* contained in the aggregate, and thus the overall aggregate size, increases with decreasing solubility. We consider iron aggregates from *k* = 1 up to *k* = 10^12^ iron ions, with corresponding radii *r_k_* ranging from 0.1 nm to approximately the size of a cell, 1 *μ*m, covering the range of iron aggregate sizes present in marine environments; in open ocean waters, iron particles radii range from less than 10nm to 350nm (von der Heyden *et al.,* 2012; Wu *et al.,* 2001). For reasons of mathematical tractability, we assume a homogeneous distribution of iron aggregates, i.e. all aggregates have the same *k*. By varying *k*, we can then isolate the effect of aggregate size on iron uptake.

### Low iron solubility strongly decreases the rate of direct iron uptake

To establish a baseline reference for the iron uptake rate, we first consider a single cell that relies on secretion-independent uptake by direct physical contact with iron, mediated, for example, by cell-attached siderophores or receptors (Fig. 1, top). The cell can only take up iron it encounters directly; in the absence of any “active” motion, this can only be mediated by random diffusion. The uptake of iron can thus be described as a set of diffusing particles in three dimensions with the cell at the origin acting as an absorbing sphere (see Methods).

**Figure 1:**
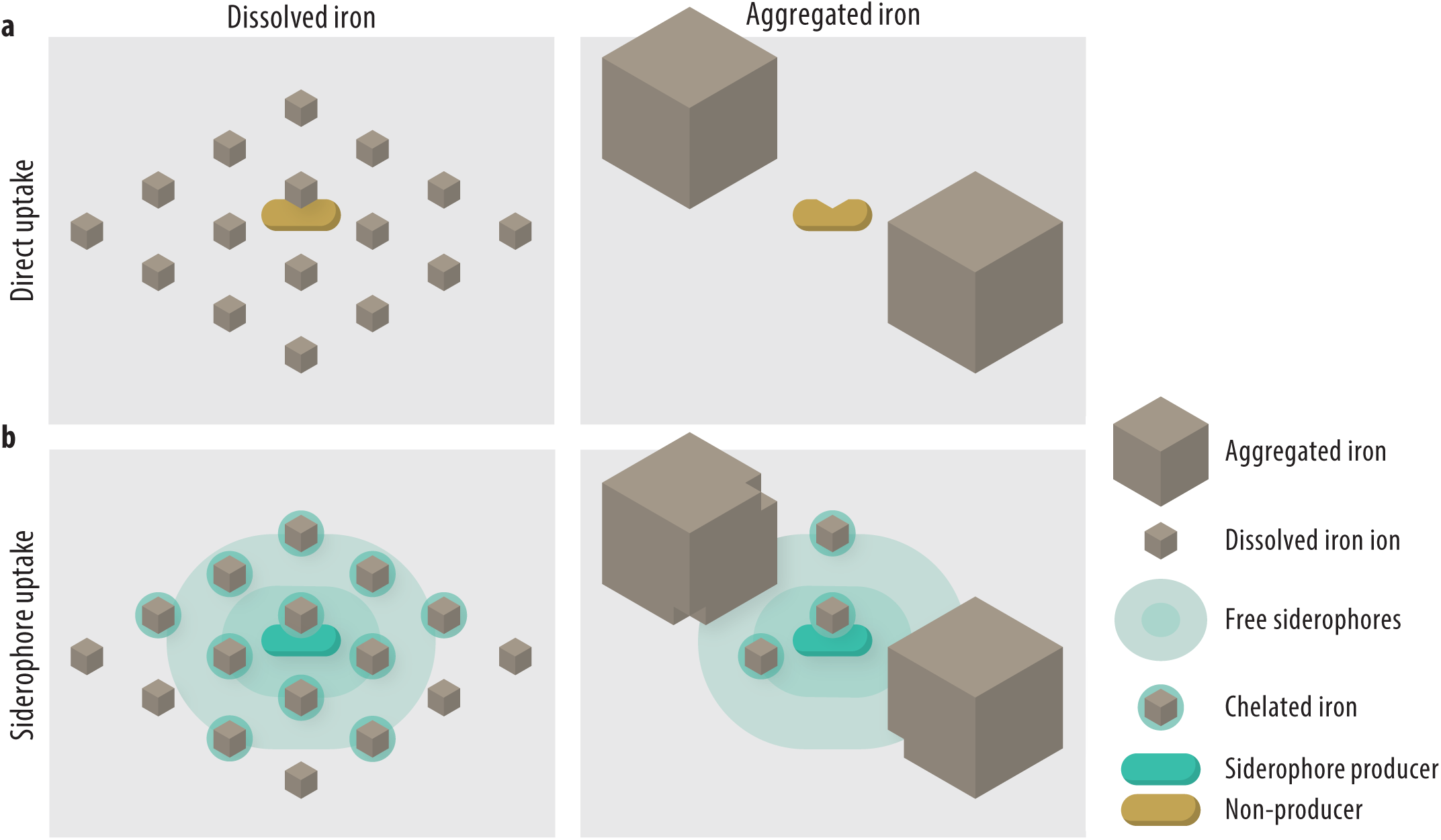
Modeling the direct and siderophore-mediated uptake of dissolved and aggregated iron. This figure illustrates two extreme cases of very strong iron aggregation or complete solubility. **a.** A cell not secreting siderophores must directly encounter iron particles for uptake. Due to its size, aggregated iron diffuses more slowly than dissolved iron. Therefore, at low solubility, which promotes iron aggregation, the cell’s iron uptake rate slows down. **b.** A siderophore-secreting cell takes up siderophore-bound iron only. Each siderophore must first chelate an iron ion before the resulting chelate can be taken up by the cell. Since chelates are larger than dissolved iron, they diffuse slower, making siderophore-mediated iron uptake slower than direct uptake. When iron is aggregated, though, the diffusion coefficient of chelates is considerably higher than that of the aggregates. Hence, use of secreted siderophores accelerates uptake at low iron solubility.

The likelihood of encounter between a cell and iron depends on the effective diffusion coefficient, which is the sum of the diffusion coefficient of the cell, *D_B_*, and of the iron, *D_k_*. The more iron ions are contained in an aggregate (larger *k*), the larger the aggregate and the smaller the diffusion coefficient, *D_k_* ~ *k*^−1/3^. As a consequence, the likelihood of encounter with the cell is smaller for larger aggregates (see Methods). Since generally *D_k_* ≫ *D*_*B*_, the uptake rate of *k*-aggregates for a single non-secreting cell with radius *R_B_* is approximately

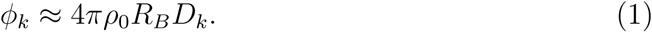

Hence, the uptake rate decreases as *k*^-1/3^ with increasing aggregate size *k*. The uptake rate also increases linearly with the background iron concentration, *ρ*_0_. We implemented a maximal uptake rate assuming an upper limit for the transport rate of iron across the cell membrane due to physiological or physical constraints (e.g. due to a maximum number of transporters; see Methods).

We distinguish between two environmental regimes, akin to diffusion-controlled reactions (Calef & Deutch, 1983): (1) In a transport-limited (reaction-limited) environment, iron uptake is limited by the amount of iron that can be processed by the transporters at the cell surface; (2) In a diffusion-limited environment, iron uptake is limited by the transport of iron to the cell surface by diffusion. Since iron aggregation influences the concentration of iron at the cell surface, the level of iron aggregation influences whether the environment is transport- or diffusion-limited at a given iron concentration (Fig. 2a).

**Figure 2:**
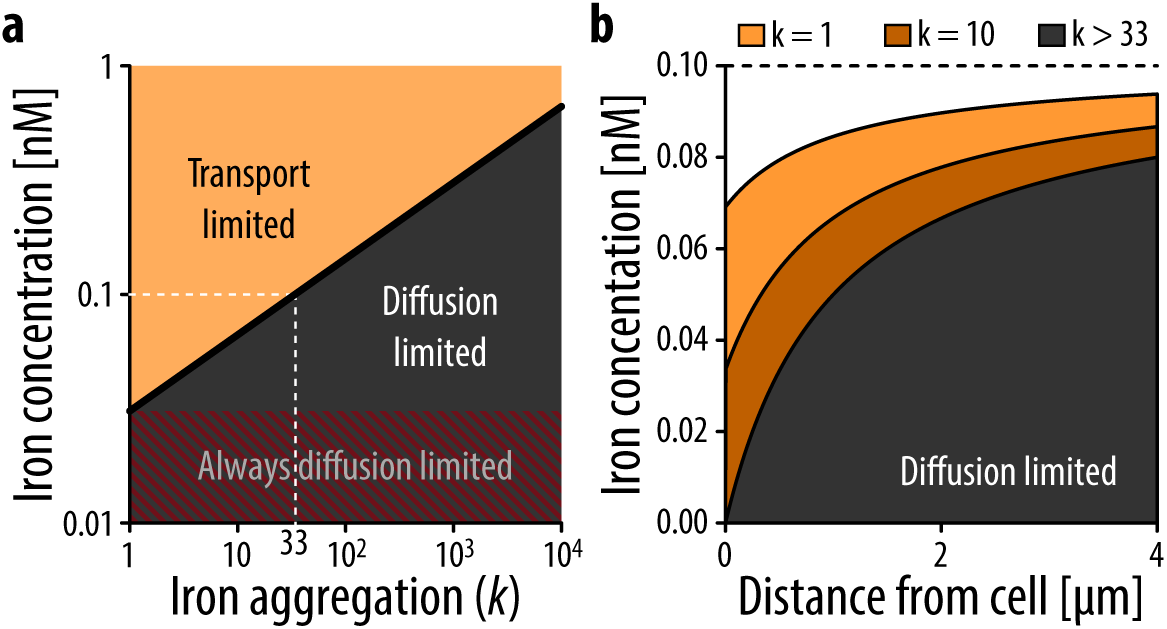
Effect of iron solubility on direct iron uptake by a single cell. **a.** Iron aggregation influences what process iron uptake is limited by. At a high iron concentration and/or if iron is highly dissolved, uptake of iron is limited by the speed of transport of iron into the cell (transport limited, orange area). As the aggregation level of iron increases at a given concentration, diffusion of iron slows down and eventually the iron at the cell surface is completely depleted. In this regime all iron that arrives at the cell surface is immediately taken up, and the cell is limited by the amount of iron diffusing to the cell (diffusion limited, black area). At very low iron concentrations, iron uptake is always limited by diffusion, irrespective of the degree of iron aggregation (red area). The dotted line highlights the background concentration used for calculations leading to the data in Figure 2b (0.1 nM). At this concentration, the threshold between the two types of limitation is at an aggregation level of *k* = 33. **b.** The equilibrium iron concentrations as a function of the distance from the cell, with a background concentration of iron *ρ*_0_ = 0.1 nM. At low *k*, in the transport-limited regime (*k* < 33, orange areas), the iron concentration at the cell surface decreases with increasing aggregation level of iron *k*, due to increasingly slow diffusion of iron, but iron is never completely depleted. In the diffusion-limited regime (*k* ≥ 33, black area), iron is completely depleted at the cell surface, and thus the steady state concentrations do not depend on the aggregation level *k* anymore. Note that the uptake rate still decreases with increasing aggregation level, even though an equilibrium concentration is reached.

At an iron concentration that is realistic for open ocean waters, *ρ*_0_ = 0.1 nM, diffusion limitation already occurs at aggregation levels of *k* ≥ 33, i.e. when iron particles have a diameter larger than only 0.3nm (Fig. 2a). We determined the equilibrium distribution of iron aggregates by solving the spherically symmetric diffusion process (see Methods and Völker & Wolf-Gladrow, 1999). The distribution of particles at steady state (Fig. 2b) illustrates that in the transport-limited regime (*k* < 33), iron is never completely depleted at the cell surface, even though the iron concentration decreases towards the cell surface. In contrast, in a diffusion-limited regime, iron is completely depleted at the cell surface and only approaches background concentration levels at a distance of over four cell radii. Under these conditions, the required time for a cell to take up sufficient iron to divide can reach up to days, for *k* ≈ 10^8^, or weeks, for *k* ≈ 10^11^ − 10^12^ (at an iron concentration of *ρ*_0_ = 0.1 nM; see Methods). Thus, a main consequence of low iron solubility is that cells can suffer from strong diffusion limitation, even at high iron concentrations.

Transport limitation can be alleviated at the cell membrane, for example through more efficient or more numerous transporters. Diffusion limitation, however, can only be overcome by influencing iron away from the cell. Secreted molecules like siderophores can modify iron sources and alter their transport properties to increase diffusion (Fig. 1, bottom). In the following, we will hence quantify the effect of siderophore secretion on overcoming diffusion limitation by increasing the diffusion speed of iron.

### Secreted siderophores can transiently increase the uptake rate of iron from large aggregates by accelerating diffusion

We next investigated whether siderophore secretion could alleviate diffusion limitation. In addition to the diffusion of iron, we accounted for free (unbound) siderophores that are produced at the cell surface and diffuse away from the cell, as well as the reaction of free siderophores with iron resulting in siderophore-iron complexes outside the cell (Fig. 1b). These complexes diffuse freely and can be taken up by the cell upon an encounter. We model these processes using a set of reaction-diffusion equations (see Methods). In order to investigate the direct effects of iron uptake mediated by secreted iron chelators, we excluded the possibility that cells take up free, unbound iron in these calculations.

Because siderophore-iron complexes are smaller than most iron aggregates, their diffusion coefficients are generally larger. This leads to accelerated diffusion that could potentially mitigate the limitation caused by the slow movement of large iron aggregates. We find that the effects of secreting siderophores depend both on the size of the iron aggregate and whether the system is still in a transient phase or has already reached steady state. In the presence of highly aggregated iron, siderophore secretion indeed results in a high uptake rate that surpasses that of direct uptake (Fig. 3, solid versus dashed lines). However, this increase is only transient: As the cell starts to produce siderophores, these solubilize the large, slowly moving aggregates close to the cell into small, fast-moving iron-siderophore chelates. This leads to an initial increase in the iron uptake rate, within the first seconds under our conditions (Fig. 3). However, the cell rapidly consumes the chelates in its proximity, and is surrounded more and more by unbound siderophores that do not encounter iron particles (Supp.Fig. S1). Therefore, as iron around the cell becomes depleted, the uptake rate begins to slowly decrease towards the equilibrium value: This decrease can take days or months, depending on the iron aggregation level (Fig. 3). At eventual equilibrium, the generation of iron-bound siderophores relies on the diffusion of large iron aggregates towards the cell. This results in the formation of a boundary layer at a distance *R*\* from the cell, where siderophores chelate unbound iron (see Supp. Fig. S1). Because this process is limited by the diffusion of iron to the boundary layer, the cell suffers from diffusion limitation equal to the case of direct iron uptake. Therefore, the equilibrium uptake rate is roughly the same as the direct uptake rate, for large enough siderophore secretion rates or low iron concentrations, *R_B_* ≫ *D_F_ρ_o_*/*P* (see Methods, Eq. 13). Overall, siderophore secretion increases the iron uptake rate compared to direct uptake, however only during a − potentially long–transient phase.

**Figure 3:**
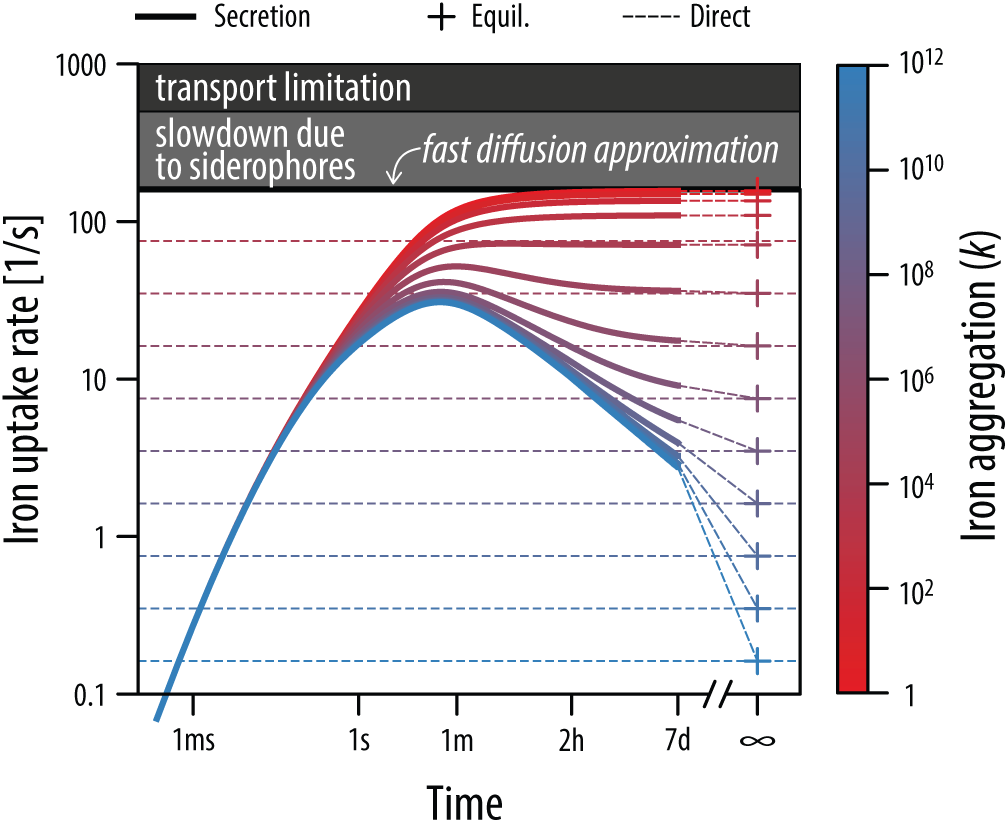
Uptake rate for secreter cells. The solid lines show the iron uptake rate for cells that secrete siderophores at a rate *P* = 0.045 amol/h. The dashed lines in the background show the corresponding direct uptake rate for reference. The uptake rate initially increases as the concentration of siderophores builds up. For low aggregation levels (reddish solid lines), the uptake rate eventually approaches the fast diffusion approximation: the approximation for the maximum uptake rate using a secretion strategy for small aggregates (bottom of grey box; see Methods). For large aggregation levels (bluish solid lines), a transient peak in the uptake rate is reached that is well above direct uptake levels, and the uptake rate only slowly approaches the equilibrium value (+ symbols). The dark grey area indicates the maximum uptake rate limited by transport into the cell.

If iron is present in small aggregates, i.e. highly soluble, the equilibrium uptake rate reached via siderophore secretion is mostly far below the direct uptake rate: releasing siderophores slows down iron acquisition. In this regime, the *fast diffusion limit*, the diffusion speed of iron is sufficiently high that the background concentration of iron can be assumed constant (see Methods, Eq. 14). Secretion slows down uptake because of two factors: First, siderophores must encounter iron and bind it, introducing an additional step prior to uptake. Second, the diffusion speed of siderophore-iron complexes is lower than that of free iron for *k*< 1000, thus reducing the flux of bound iron towards the cell. For small iron aggregates (highly soluble iron), siderophore secretion thus actually slows down uptake. Therefore, for a range of *k* = 33 to *k* = 1000 (at an iron concentration of 0.1 nM), the cell is diffusion-limited, but secretion of siderophores is not effective at overcoming this limitation.

To access poorly soluble iron, cells could also, instead of secreting siderophores, increase their own diffusion coefficient by engaging in swimming motility. However, to achieve sufficient iron uptake, the cell would need to swim at a significant speed: Already for moderate iron aggregation levels of *k* > 10^5^, the cell would need to increase its diffusion coefficient over 100-fold to reach an effect comparable to siderophore secretion at a rate of 1amol/s upwards (see Supplementary Information for more detailed discussion). While this is within the range of measured swimming speeds, especially for marine bacteria (Stocker & Seymour, 2012), swimming also implies large metabolic costs (Mitchell, 2002; Taylor & Stocker, 2012), and this is likely exasperated in nutrient poor environments. Our model calculations show that siderophore secretion is an alternative to active motility.

### The effect of siderophore secretion depends on siderophore production rate and iron aggregation

In the previous section we showed that siderophore secretion can transiently speed up iron acquisition. To investigate how these transient effects scale with bacterial growth, we define the ‘uptake time’ as the required time until a cell has acquired enough iron to produce enough biomass to initiate cell division (assuming an iron content of 10^6^ Fe atoms = 1.66amol per cell (Andrews *et al.,* 2003); Fig. 4a). Note that, if cells are limited by iron only, the uptake time is equal to the inter-division time.

**Figure 4:**
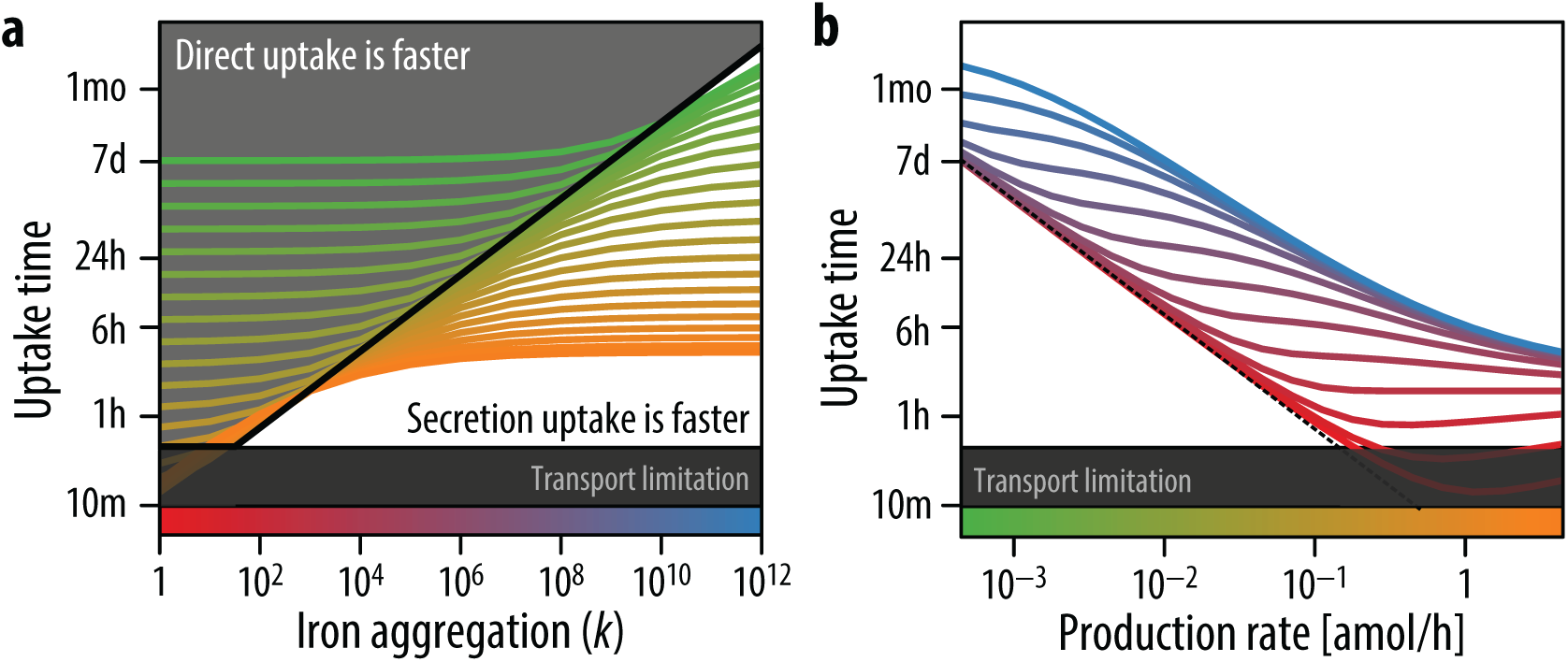
Uptake time for secreter cells. **a.** The time required to take up enough iron for division (‘uptake time’) depends on the iron aggregation level and the siderophore production rate. The black line indicates the uptake time for a cell relying on direct uptake. Aggregation level and production rate combinations that fall to the left of this line are values where secretion-based uptake is slower than direct uptake (grey area). Aggregation level and production rate combinations that fall to the right are conditions where secretion-mediated uptake is faster (white area). For low iron aggregation levels, a secreter has higher uptake times than a non-secreter, unless it produces siderophores at a very high rate (> 1amol/h). For large iron aggregation levels, a secreter shortens its uptake time compared to a non-secreter, even for low production rates (< 0.01amol/h). The full color scale for the production rate is shown in Panel b on the x-axis. **b.** Uptake time is influenced by the siderophore production rate. For highly dissolved iron (more red lines) or very low production rates, this relationship is almost linear. For highly aggregated iron (more blue lines), the effect is slightly reduced (blue lines not as steep). For low aggregation levels, or low levels of production for high aggregation levels (not visible), the uptake time is well approximated by the fast diffusion approximation (dashed line). As the production rate increases from very low levels, the relationship between production rate and uptake time flattens out, indicating that an increase in production rate only has a small effect on reducing the uptake time. As the production rate increases further, the effect on reducing uptake time becomes stronger again. The flattest range corresponds to where the secretion and direct uptake strategies result in approximately the same uptake time. The full color scale for the aggregation level is shown in Panel a on the x-axis.

The increased iron uptake rate achieved by secretion at low iron solubility, although only transient, can significantly shorten the uptake time (Fig. 4a). A non-motile cell relying on direct physical contact has an uptake time of around 15 days for parameters close to the marine environment (iron concentration of *ρ*_0_ = 0.1 nM; aggregation level of *k* = 10^10^, i.e. aggregate radius of around 215nm). When a cell starts to secrete siderophores, the uptake time at these conditions is significantly reduced to a few days or hours, depending on the production rate. Conversely, for low aggregation levels, i.e. highly soluble iron, a secreter cell has a longer uptake time than a cell using direct uptake. The uptake time is strongly influenced by the production rate of siderophores. Overall, the higher the siderophore production rate, the wider the range of environments in which siderophore secretion speeds up iron acquisition compared to direct uptake. At low *k* and low siderophore production rates, the uptake time decreases linearly with the production rate (slope −1 of red line in Fig. 4b). Hence, a two-fold increase in the secretion rate of siderophores leads to roughly a two-fold decrease in the uptake time. For large aggregation levels (high *k*), the uptake time generally decreases close to, but less than linearly with production rate (blue line, Fig. 4b). Thus, an increase in production rate still has a positive effect on the uptake time, though the return is diminished. For intermediate aggregation states, the system transitions from having non-zero concentration of unbound iron to a depletion of iron in the close proximity of the cell. In this regime, an increase in production rate has almost no effect on the uptake time (flattening of the lines in Fig. 4b). Such parameter combinations of aggregation level and production rate are also where the uptake time of both strategies is roughly similar. The diminished returns in these regimes indicate that the secreter needs to drastically increase its production rate to gain marginal benefits over a direct uptake strategy.

The cost of producing such a high number of siderophores is difficult to estimate, since the negative effect of siderophore production on growth, due to resources spent on production instead of cell division, likely depends on environmental parameters such as the level of resource limitation (Brockhurst *et al.,* 2008). However, to obtain an estimate of the magnitude and efficiency of siderophore production, we calculated how many siderophore molecules need to be produced in order to take up one iron ion (a similar approach to (Völker & Wolf-Gladrow, 1999)). For a low siderophore secretion rate of 0.045 amol/h and an aggregation level of *k* = 10^10^, a secreter must release around 10 siderophores to take up one iron ion. However, this secretion rate leads to relatively slow uptake with an uptake time of around 15 days at an iron concentration of *ρ*_0_ = 0.1 nM. Faster uptake is achieved through higher siderophore secretion rates, though this also implies that more siderophores per iron are secreted: a secretion rate of 4.5 amol/h results in a division time of 45 hours, requiring around 120 secreted siderophores per iron taken up at an iron concentration of *ρ*_0_ = 0.1 nM and for *k* = 10^10^.

### Synergistic siderophore production between two neighboring cells can occur over a wide range of parameters

High siderophore production rates probably require a substantial investment of cellular resources. One way to reduce the production effort is cooperative production of siderophores in groups of cells. Siderophore-iron chelates that diffuse from one producer cell might still benefit a producing neighbor cell. Furthermore, if the cells share their siderophores, each cell might have to produce fewer siderophore molecules. The outcome of such interactions is not easily predictable, though, since each cell is both a source of siderophores and also a sink of iron-bound chelates.

Siderophore production in a group of cells has been shown to increase the efficiency of accessing iron-bound siderophores thanks to an accumulation of siderophores in the neighborhood of the cells (Völker & Wolf-Gladrow, 1999). However, these results did not consider the effect of poorly soluble iron aggregates. In our model we observe that, especially in the case of poorly soluble iron, siderophore production affects and alters the environment in a large neighborhood around the cell. Iron-bound siderophore chelates build up at distances of several hundred *μ*m (see Supp. Fig. S1), thus potentially allowing for cooperative interactions between cells located at significant distances. Our model indicates that both costs and benefits are influenced by the solubility of iron: First, at low iron solubility fast uptake from large iron aggregates entails a high siderophore-to-iron expenditure ratio, indicating that an important proportion of siderophores is lost to the producer cell even if no other cell is present. Thus, direct of costs of sharing may be minimal at low solubility. Second, the benefits of sharing siderophore production with a producing neighbor cell are influenced by the beneficial returns of increasing siderophore production rate (Fig. 4b).

We investigated a first stage of growth in a group of cells by considering the interactions between two cells. We measured the average number of siderophores produced by a secreter cell per iron taken up during the uptake time. The net effect of growing in proximity to another cell depends on how much a cell suffers from competition for iron-bound siderophores compared to how much it benefits from the siderophores produced by its neighbor. If sharing siderophores is synergistic, i.e. more efficient in a group, a cell has to produce fewer siderophores to take up one iron ion when a producing neighbor is present. We first quantified how much a siderophore-secreting cell is negatively influenced by the presence of a second cell by implementing this cell as a iron-siderophore chelate sink (comparable to cheater cells in Griffin *et al.,* 2004). We assume that the second cell does not produce siderophores, but still can take up iron bound to siderophores (but not free iron). We measure the negative effect (or cost) in terms of how many more siderophores the secreter cell needs to produce on average to take up one iron, relative to a situation where no other cell were present. A value of 1 indicates that the production effort of the producer is not altered by the presence of the nonproducing cell, whereas a value of e.g. 2 means that twice as many siderophores need to be produced to take up one iron. We find that, while the degree of iron aggregation does not strongly influence the interaction, the distance between the cells plays a key role (Fig. 5c). If the two cells were (hypothetically) at the exact same point in space, then all iron-siderophore chelates that arrive at the cells are shared evenly between the two, and the producing cell needs to produce approximately the double amount of siderophores (only approximately because the uptake rate is not linear in time, see Fig. 4b). As the distance between the two cells increases, the negative effect on the producer decreases (Fig. 5c). For a secretion rate of 4.5amol/h, the loss of siderophores to the neighboring cell at distances larger than 10 *μ*m has a negligible effect on the producing cell (Supp. Fig. S5). At the same time, at these distances, the nonproducing cell is able to take up almost equivalent levels of iron as the secreting cell (Fig. 5d). Thus, siderophores can efficiently be taken up by the neighboring cell at a very low cost to the producer, because the non-producer benefits from the siderophores that would likely be lost to the producer anyway. Iron aggregation has a strong influence only in one aspect: The nonproducing cell, even if located at large distances from the producer, can still benefit from the siderophores, however only if iron is aggregated. If iron is highly dissolved, most chelates are produced close to the cell (Supp. Fig. S1) and are rapidly taken up by the producer, such that the nonproducing cell is only able to take up a fraction of the iron relative to the producing cell (Fig. 5d).

**Figure 5:**
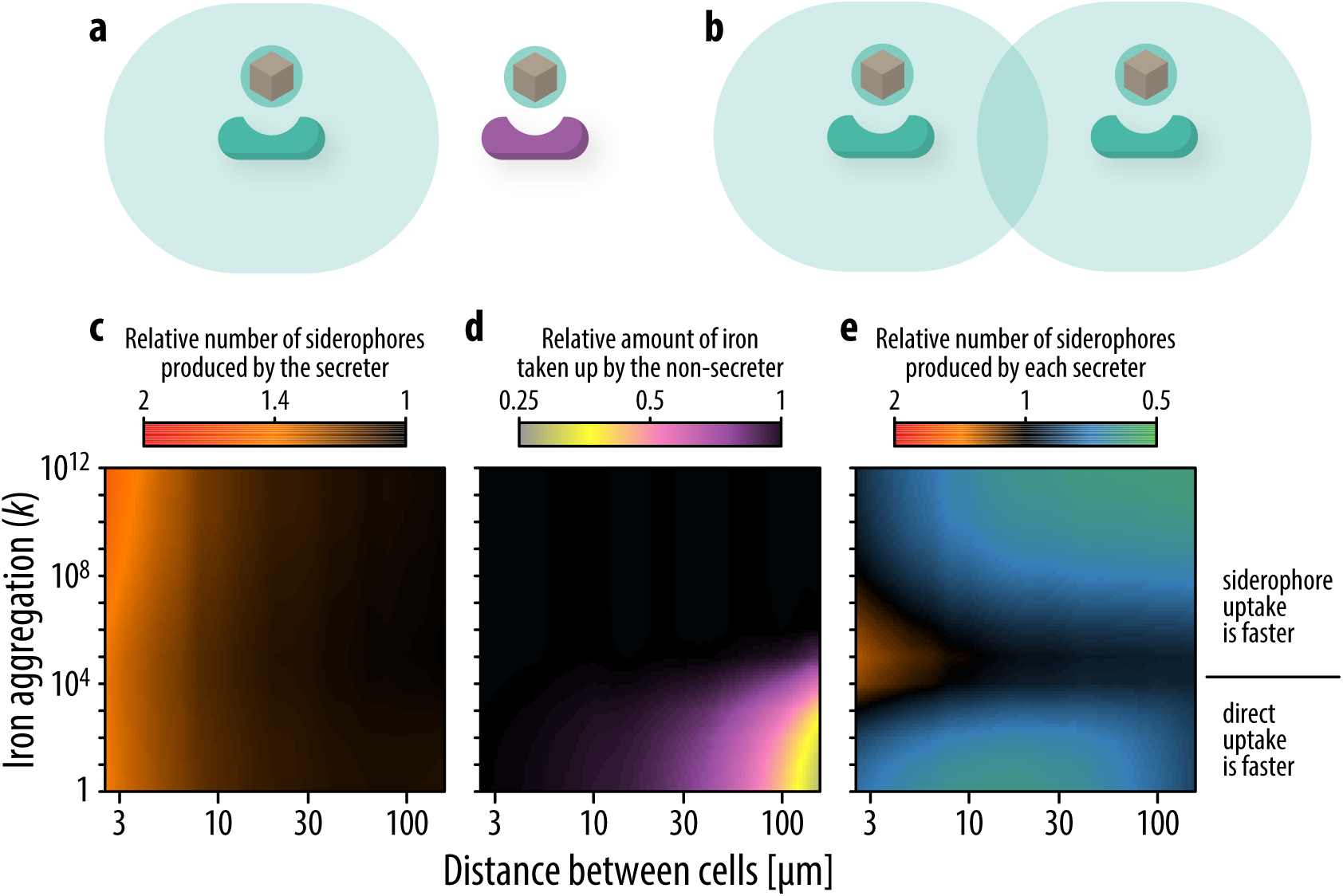
Effect of two competing cells on each other. a,c,d. Only the secreter produces siderophores, the non-secreter consumes iron-bound siderophores, but does not contribute to production. **c.** The relative increase in the average number of siderophores that need to be produced to take up one iron, relative to if the secreter was alone, depends on the distance between the cells. When the two cells are close, the amount of siderophores required increases almost twofold (orange area). However, when cells are at distances of >30 *μ*m, the secreter does not need to produce additional siderophores (black area). **d.** The relative amount of iron acquired by the non-secreter in the time span the secreter takes up one iron ion is influenced by the cells’ distance. The non-secreter acquires less iron relative to the secreter only for low aggregation levels and at large distances (purple/yellow area). **b,e** Both cells produce siderophores. **e.** Secreters can increase the uptake rate of a neighboring secreter at no or little additional cost, provided that the distance between cells is large enough. For low iron aggregation levels, this reduces the slowdown due to secretion compared to direct uptake (lower blue/green area). For high aggregation levels, this increases the acceleration of uptake by secreting siderophores. At intermediate aggregation levels the marginal benefit of producing additional siderophores is small, though, leading to no benefits or even net costs (black and orange area, respectively).

Since the benefits of siderophore secretion can be shared with neighboring cells at relatively short distances without additional costs to the producing cell, we next examined how joint secretion may be synergistically beneficial for cells. We measure benefit here as a reduction in the amount of siderophores secreted to achieve uptake of one iron ion. If this value is 1, then there is no net effect of the presence of another cell, and competitive and cooperative effects balance each other. A value larger than 1 indicates that there is a net negative effect due to competition, i.e. cells suffer from their neighbors taking up iron-bound siderophores. On the other hand, a value smaller than 1 means that both cells benefit from the presence of another producer, because they efficiently share siderophores. As a result, each cell has to secrete fewer siderophores to take up the same amount of iron. Over a large range of separation distances (> 10*μ*m for *P* = 4.5amol/h), cells can share the benefits of secretion without an additional cost due to competition (blue-green areas in Fig. 5e), and cooperative effects are thus greater than competitive effects. The magnitude of the synergistic effect, however, depends on the marginal benefit of increased siderophore production rate, i.e. the slope in Fig. 4b. For production levels that only result in a small increase in iron uptake, the marginal benefit of an additional producer cell is small (black areas in Fig. 5e), which is the case for iron aggregates of intermediate size. At other conditions, however, there is a strong effect and the amount of siderophores produced for the uptake of 1 iron can be reduced almost twofold.

The outcome of social interactions is therefore strongly influenced by the physical properties of iron, in particular the diffusion coefficient. Highly soluble and insoluble iron sources enable synergistic interactions, whereas medium-sized aggregates do not. Overall, we find that at the majority of our conditions, siderophore secretion can lead to synergistic interactions between two neighboring siderophore producing cells. This aids in making the secretion of siderophores a favorable strategy in a wide range of environments compared to direct uptake (Fig. 5d).

## Discussion

The ubiquity of bacterial siderophore secretion despite the risk of siderophore loss and the evolutionary fragility it entails has stimulated a large body of research (Griffin *et al.,* 2004; Kümmerli & Brown, 2010; Dumas & Kümmerli, 2012). Here, we suggest that the release of siderophores is essential for their function, potentially explaining why this uptake strategy is so widespread. We show that low iron solubility, present in many environments, can strongly slow down iron uptake if no siderophores are released. Secreted siderophores solubilize iron and generate small chelates with significantly increased diffusion speed. However, our results show that secretion of siderophores only increases iron acquisition rates when iron aggregates are large, and is generally not beneficial at high solubility, when aggregates are small.

In vertebrate hosts, iron is not aggregated, but solubilized by chelating proteins, e.g. transferrin or lactoferrin (Baker & Baker, 2005; Gomme *et al.,* 2005). Our results suggest that such environments containing solubilized iron should not be conducive to siderophore-based iron uptake. However, the chelating host proteins are considerably larger than siderophores, such that siderophores would still accelerate the diffusion of iron: Transferrin and lactoferrin have a molar mass of around 79 kilodaltons, whereas the molar mass of siderophores ranges from around 0.1–1.4 kilodaltons (Kümmerli *et al.,* 2014). The magnitude of this acceleration will likely be smaller than in environments with poorly soluble iron, potentially explaining why some pathogenic bacteria use receptors for direct uptake of iron-chelate complexes instead of siderophores (Braun & Killmann, 1999) and why pathogens can lose siderophore systems during adaptation to the host environment (Andersen *et al.,* 2015; Marvig *et al.,* 2014).

In our model we have made two assumptions that potentially strongly disfavour a siderophore secretion strategy. First, secreter cells cannot take up iron through direct physical contact. It is likely, however, that siderophore-producing cells are able to take up iron that is directly encountered, since bacteria possess several iron uptake systems (Andrews *et al.,* 2003; Wandersman & Delepelaire, 2004). In this case, a siderophore producer can take up fast diffusing small iron aggregates as well, mitigating the negative effects of acquisition slowdown at high iron solubility. Second, we assume that large iron aggregates are immediately taken up by a non-producing cell upon physical encounter. Realistically, cells would need to process the iron aggregates, thus slowing down the direct uptake of large aggregates. By using siderophores, cells can “pre-process” this iron for fast uptake at a distance from the cell. Since both of these assumptions enhance the benefits of direct uptake we measure in our results, the environments predicted by our model to be conducive to siderophore secretion are likely conservative estimates.

Siderophore secretion is often cited as one of the central examples for diffusible public goods in the study of the evolution of cooperation (Velicer, 2003; Griffin *et al.,* 2004; Kümmerli *et al.,* 2009; Allen *et al.,* 2013). Secreted molecules, such as siderophores, are at risk of being “stolen” by a cheater genotype that avoids the cost of production, but reaps the benefits. The current consensus is that cooperation by means of secreted public goods is stabilized by spatially structured environments, as this increases the probability of interactions between identical genotypes, and consequentially decreases the probability of interactions between producers and cheats (Kümmerli *et al.,* 2009; Nadell *et al.,* 2010; Julou *et al.,* 2013; Allen *et al.,* 2013). Only few of these studies, however, have considered the process of diffusion of the public good to the full extent (Vetter *et al.,* 1998; Allison, 2005; Folse & Allison, 2012; Allen *et al.,* 2013; Dobay *et al.,* 2014). More importantly, game-theoretical studies mostly model the diffusive public good itself as a carrier of some benefit (Allen *et al.,* 2013; Dobay *et al.,* 2014). In our study, we consider a more realistic view of the mechanism of siderophore-based iron uptake. A secreted siderophore *per se* is useless to the cell, and any neighbouring cell, until it encounters and chelates iron. A detailed understanding of how the benefit of siderophores, efficient iron uptake, is generated, emphasizes the importance of diffusion of siderophores. At low iron solubility, secreted siderophores are a means to overcome diffusion limitation, and hence the diffusion away from the producer cell is part of the siderophore’s function. Therefore, reduced diffusivity of siderophores (Martinez *et al.,* 2003; Kümmerli *et al.,* 2014; Scholz & Greenberg, 2015) can likely stabilize siderophore production in the face of the public goods dilemma, but it probably also reduces the efficiency with which siderophores improve iron uptake at low iron solubility.

The location of where benefits of siderophores are generated is the result of a complex interplay between iron sources and the producer’s location. This is likely to have an effect on the interactions between all types of cells. Therefore, abiotic properties of the environment, in this case the diffusivity of iron, can play an important role in social interactions between cells, and need be taken into account when considering cooperative dynamics. The same also applies to other secreted metabolites that interact with large slow substrates, such as chitinases secreted to degrade chitin, or other extracellular degrading enzymes (cellulase, exoprotease).

We suggest that when analyzing microbial competition and cooperation it is important to consider two different length scales: (1) the length of competition, i.e. the distance between neighboring producers at which they compete for the shared resource, in this case the iron-bound siderophores; and (2) the length of synergism, i. e. the distance at which the neighboring producers synergistically utilize the siderophores produced. Our model shows that typically the length scale of competition is much shorter than the length scale of synergism. This allows neighbouring producer cells to jointly increase the global siderophore production without paying additional costs due to competition. Thus, the benefits of siderophore secretion increase with the number of cells, while the costs per cell stay constant, if the cells are sufficiently spaced. This could set the basis for successul cooperative interactions in the secretion of many compounds.

## Methods

### Direct uptake

In the direct uptake case, iron is taken up when a cell encounters an iron aggregate. If we assume a spherical cell with radius *r_B_*, then the probability that a spherical iron aggregate starting at a distance *r* from the cell encounters the cell before time *t* is just the hitting probability of a random walk (Crank, 1975; Frazier & Alber, 2012),

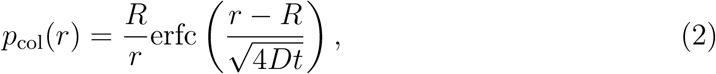

with total radius *R* = *r_B_* + *r*_Fe_ and effective diffusion coefficient *D* = *D_B_* + *D*_Fe_. The diffusion coefficient is, *D* = *k_B_T*/6*πhr*, where *k_B_* ≈ 1.38 × 10^−23^ JK^−1^ is Boltzmann’s constant, *T* = 293 K is ambient temperature, *h* = 1.003 mPa.s is the viscosity of water at ambient temperature, and *r* is the radius of the spherical particle in meters. By summing the diffusion coefficients to an effective diffusion coefficient, we fix the reference frame of the iron particle to the bacterium and subsume the movement of the bacterium into the movement of the iron particle. As a simplification, we assume that the cell is stationary in space, and thus the effective diffusion coefficient is *D* ≈ *D*_Fe_. Note that generally iron diffusion is faster than the cell, *r_B_* ≫ *r*_Fe_, so that in most cases *D_B_* + *D*_Fe_ ≈ *D*_Fe_ is an acceptable approximation.

The total number of particles that have collided with the bacterium by time *t* is,

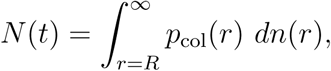

where *dn*(*r*) is the number of particles at a distance *r*. If the concentration of iron is *ρ* and the iron particles are equally distributed in space, then *dn*(*r*) = *ρ* · 4 *πr*^2^ *dr*. Thus the total number of particles becomes,

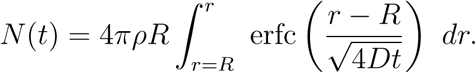

We assume that all iron particles are aggregates of size *k* and that the total concentration of iron is *ρ*_0_ = *k* · *ρ*_*k*_, where *ρ*_*k*_ is the concentration of particles of size *k*. The number of particles of size *k* that have collided with the bacterium at time *t* is,

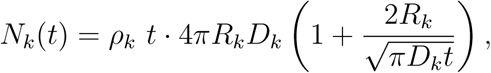

with *R_k_* = (*r_B_* + *k*^1/3^*r*_Fe_) and *D_k_* = *k_B_T*/(6*πh* · *k*^1/3^*r*_Fe_). At best, a bacterium can completely take up a *k*-aggregate, such that the total amount of absorbed iron is,

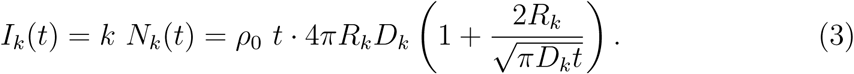

Note that the assumption that bacteria can take up complete iron aggregates is equivalent to assuming that the iron is fully dissolved, but that the individual atoms diffuse with a reduced diffusion coefficient *D_k_*. In this case, the concentration of iron, *F*(*r*, *t*), can be equivalently represented as a spherically symmetrical diffusion process,

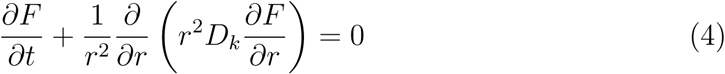

with an absorbing boundary at *r* = *R_k_*, a reservoir at infinity, lim_*r*→∞_*F*(*r*,*t*) = *ρ*_0_, and initial concentration *F*(*r*, 0) = *ρ*_0_. The general equilibrium solution to Eq. 4 with 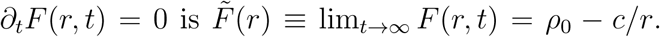 The constant *c* is found by the boundary condition at *r* = *R_k_*. On the one hand we require *F*(*R_k_*,*t*) ≥ 0, and thus *c* ≼*R_k_ρ*_0_. On the other hand the iron uptake is limited by the maximal transport rate. The flux across the cell surface is,

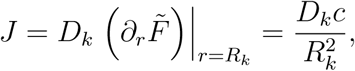

and thus the maximal flux for *c*^max^ = *R*_*k*_*ρ*_0_ is *J*^max^ = *D_k_ρ*_0_/*R_k_*, the same as the asymptotic limit of Eq. 3. We say that the cell is iron-diffusion limited if the maximal flux is smaller than the maximally possible transport rate, i.e. *J*^max^ < *α* (see Fig. 2b).

### Siderophore-mediated uptake

The iron uptake by a single secreting cell is modelled as a reaction-diffusion process for free iron, *F*, unbound siderophores, *X*, and bound siderophores, *Y*,

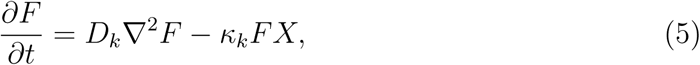

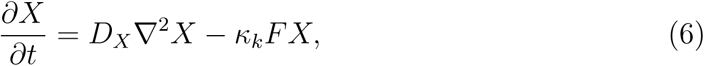

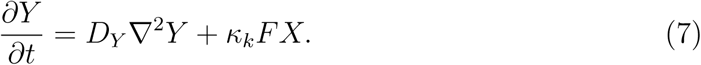

When we only consider a single cell, the system is spherically symmetric. Thus the reaction-diffusion equations become (see also Völker & Wolf-Gladrow, 1999),

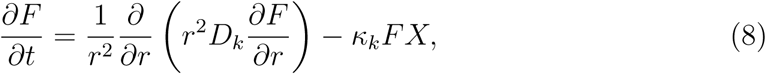

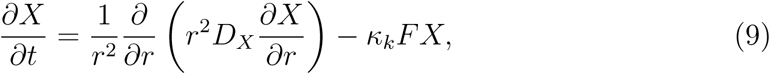

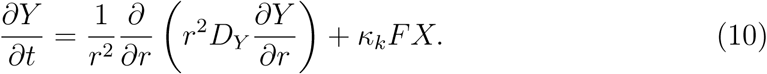

We assume that the binding affinity is the same for all aggregation levels, *K_k_* ≡ *k* = 10^6^ M^-1^s^-1^ and that the diffusion coefficient of free siderophores the same as for iron-siderophore complexes. Initially, iron is equally distributed in space at a concentration of *F*(*r*, 0) = *ρ*_0_ and there are no siderophores present, *X*(*r*, 0) = 0. We assume that the concentration of free and bound siderophores tends to zero far away from the cell, lim_*r*→∞_ *X*(*r*, *t*) = *Y*(*r*,*t*) = 0, and the concentration of iron is constant far away from the cell, lim_*r*→∞_ *F*(*r*,*t*) = *ρ*_0_. The uptake rate of iron-siderophore complexes by the cell is then just equal to the flux of *Y* at *r* = *R_B_*,

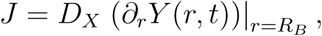

with a maximal rate of *α* as in the direct uptake case.

To gain some analytical understanding of the equilibrium distributions, we consider some limiting cases of the reaction-diffusion system.

#### No reaction

In absence of any reaction of siderophores with iron, *κ* = 0. In this case the equilibrium solution for the distribution of free siderophores is,

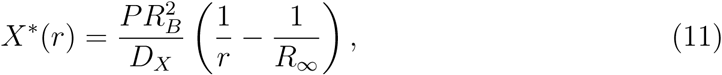

where *P* is the excretion rate of siderophores from the cell. Here, *R*_∞_ is the upper bound of the considered volume. For an open system, *R*_∞_→∞.

#### Large aggregation

When the level of aggregation is large, then the diffusion of free iron is much slower than the diffusion of free siderophores. But since the concentration of siderophores decreases with distance from the cell as a consequence of the spherical dilution, there exists a boundary at *r* = *R*^*^, where the influx of free siderophores completely reacts with the influx of free iron.

Let ∆*X* = *ϕ_x_* ∆*t* and ∆*F* = *ϕ_F_* ∆*t* be the amount of *X* and *F* that enters a finite small volume during time ∆*t*. Then the amount that reacts will be ∆*y* = *k*∆*F*∆*X*. We are interested in the case where the all iron ∆*F* reacts with all siderophores ∆*X*, *k*∆*X*∆*F* = ∆*F* and *k*∆*X*∆*F* = ∆*X*, and hence, ∆*F* = ∆*X*, or, |*ϕ_x_*(*R*^*^)| = |*ϕ_F_* (*R*^*^)|.

Iron diffuses to *r* =*R*^*^ from above, and siderophores diffuse to *r* = *R*^*^ from below. Thus for *r* < *R*^*^, the distribution of free siderophores follows Equation (11) with *R*_∞_ = *R*^*^. Equivalently, the distribution of iron for *r* > *R*^*^ follows that of freely diffusing iron, *F*(*r*) = *ρ*_0_ (1 − *R*^*^/*r*). The fluxes are then,

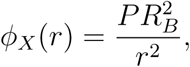

and,

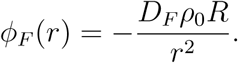

These are equal at,

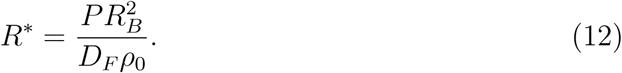

This defines a boundary at a distance *r* = *R*^*^ from the cell. Below this radius there are enough siderophores to bind all free iron and thus there is no free iron. Above this radius, all the siderophores have been bound. Hence at equilibrium, siderophore-iron complexes are only produced at this radius.

The distribution of siderophore-iron complexes above *r* = *R*^*^ then is,

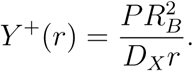

Below 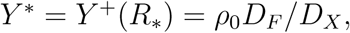,where 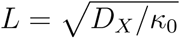,

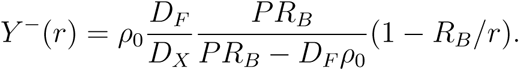

Finally, this results in a flux at the cell of,

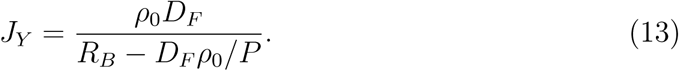

This converges to the maximal direct uptake flux, *J* = *ρ*_0_*D_F_*/*R_B_*, for secretion rates *p* ≫ *D_F_ ρ*_0_.

#### Fast diffusion

If the diffusion speed of free iron is fast compared to the reaction speed of siderophores, then the background concentration of free iron can be assumed constant. The reaction-diffusion equations then become,

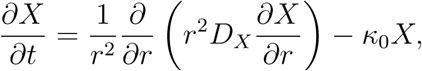

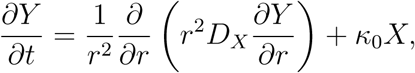

with *k_0_* = *kρ*_0_. The equilibrium solutions to these equations for boundary conditions lim_*r*→∞_ *X*(*r*) = lim_*r*→∞_ = 0, *ϕx*(*R_B_*) = −*D_x_∂_r_X*(*r* = *R_B_*) = *P* and *Y*(*R_B_*) = 0 is (see Völker & Wolf-Gladrow, 1999),

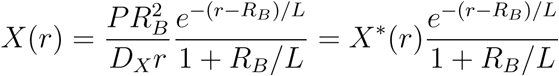

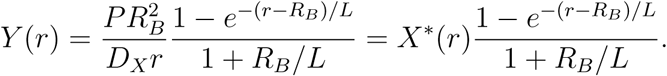

Here, 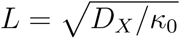 is the characteristic diffusion-reaction length of siderophores with the background iron. The derivative of *Y* is,

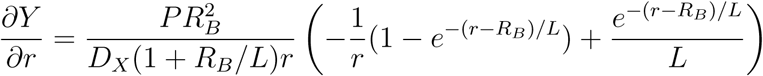

Hence, as derived in (Völker & Wolf-Gladrow, 1999), the maximum iron uptake rate is,

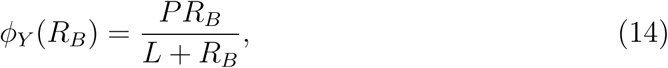

and the peak in the distribution is at a distance *r*, where

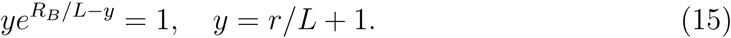

### Numerical integration of the partial differential equations

With exception of the single cell direct uptake case, no analytical solutions to the reaction-diffusion equations are available. We therefore numerically integrated the equations using a finite elements approach implemented in the FEniCS project version 1. 6 (Logg *et al.,* 2012). The FEniCS software suite uses the variational formulation of the PDEs on meshes.

#### Single cell

For the single cell case, we exploited the spherical symmetry and used a linear expanding mesh between *R_B_* = 1 *μ*m and *R*_∞_ = 10 m, and *m* = 400 mesh intervals,

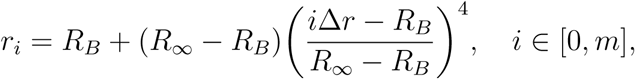

where ∆*r* = ( *R*_∞_ − *R_B_*)/m. We solved the PDEs over discrete increasing time steps, with ∆*t*_0_ = 10^−4^ s and ∆*t*_*j*+1_ = 1.2∆*t_j_*, with a maximum time step of 1000 s (see also Supplementary Information).

#### Two cells

In the two cell case, the system only has cylindrical symmetric along the axis that connects the two cells,

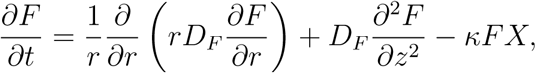

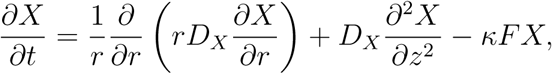

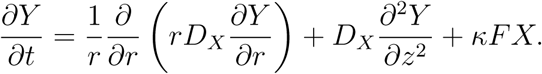

We generated 2D (*z*,*r*)-meshes using the following procedure: We first created a circular domain of radius *ρ*_1_ = 100 *μ*m. We then removed two circular ‘cells’, with radius *R_B_* and varying distance *d* from each other, from the domain. The domain was then converted to a mesh using FEniCS with a mesh size of 10, and subsequently refining all mesh elements within the circle *ρ* = the mesh was finally expanded to a full r*ρ*_1_/2. The mesh was finally expanded to a full radius of *ρ*_∞_ = 0.1 m, by adding concentric circles of *m*_2_ = 24 mesh points at increasing radii *ρ*_*j*+1_ = *ρ_j_* (1 + 2*π*/*m*_2_). Finally, we integrated the 2D reaction-diffusion equations using an increasing time step, ∆*t*_0_ = 10^−2^ s and ∆*t*_*j*+1_ = 1.2∆*t_j_*, with a maximum time step of 1000 s.

## Acknowledgements

We thank Thierry Sollberger for help with designing Figures 1 and 5 as well as Roman Stocker, Laura Sigg and Stephan Kraemer for helpful discussions. KTS and MA were supported by Eawag and ETH Zurich. GEL and KTS were supported by the Swiss National Science Foundation.

## A Variational formulation of the reaction-diffusion problem

We numerically integrate the reaction-diffusion equations for the two cell case using FEniCS 1.6.0 (Alnæs *et al.,* 2015). In order to solve the time dependent problem, we use a finite differences approximation for the time dimension, and solve the variational formulation at each time step using the finite element method. For completeness, we here briefly reiterate the formulation of the time-dependent variational formulation.

### Variational formulation for the stationary problem

The finite element method uses the weak variational formulation of the system of partial differential equations of the reaction-diffusion system. Briefly, a PDE for the function *u*(*r*,*z*) with cylindrical symmetry defined on (*r*, *z*) ∈ Ω of the form,

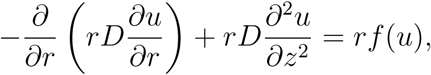

is multiplied by a test function *u*(*r*,*z*) and integrated over Ω,

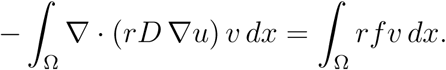

The left hand side can be integrated by parts, and by forcing *v* = 0 on the boundaries where *u* is known. The cylindrically symmetrical problem stated in the weak variational formalism is thus,

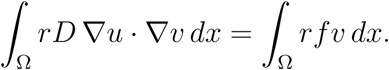

Equivalently, for a spherically symmetric problem,

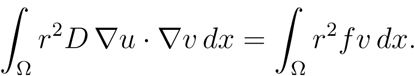

### Variational formulation time-dependent problem

Here we use the standard finite difference discretization of the time derivative, such that *∂_t_u*≈ (*u*^(*k*)^−*u*^(*k*−1)^/∆*t* The time dependent PDE equation becomes,

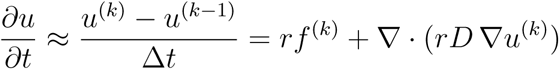

We then iterate along the finite differences by successively solving the following equation using finite elements for a known *u*^(*k*−1)^,

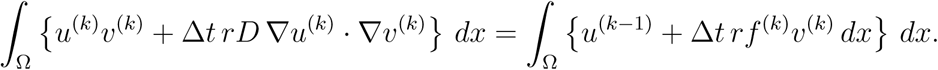

## B The spatio-temporal effects of siderophore secretion on the distribution of iron in space

Strong temporal dynamics become visible when quantifying the radial distributions of iron, siderophores and bound iron over time for aggregates of different sizes in the presence of secreted siderophores (Fig. S1). As soon as the cell begins to secrete siderophores, the iron close to the cell is quickly bound and depleted, and the iron-siderophore complexes are subsequently taken up by the cell with a high probability. If iron is soluble (*k* = 1) then the diffusion speed is maximal and the bound free iron close to the cell is quickly replenished by new iron diffusing towards the cell from a distance. Therefore, the iron concentration close to the cell is roughly the same as the background level (Fig. S1a). New siderophores thus quickly encounter free iron to bind, which can then be taken up by the cell, resulting in a high concentration of bound iron-siderophore complexes close to the cell (see Methods, Eq. 15). The cell is thus transport-and not diffusion-limited. In this fast diffusion limit the background concentration of iron is almost constant, and the iron uptake rate is primarily determined by the secretion rate of siderophores (see Methods and Völker & Wolf-Gladrow, 1999).

At higher aggregation levels, though, fresh iron diffuses towards the cell more slowly, eventually resulting in a total depletion of free iron close to the cell (Fig. S1b,c). At the same time, free siderophores diffuse away from the cell and a ‘hot-spot’ for the formation of bound siderophores builds up at a certain distance from the cell, resulting in a traveling peak in the distribution of bound iron over time (Fig. S1b,c). As bound iron is either taken up by the cell or diffuses away, this peak eventually flattens out over time and moves away from the cell to a final distance *R*^*^ (dashed lines in Fig. S1; see Methods, Eq. 12). The distribution of bound iron thus approaches a lower equilibrium level. Within this boundary, *r* < *R*^*^, the concentration of free iron is zero, i.e. close to the cell all nonchelated iron is completely depleted. At the boundary, *r* = *R*^*^, the influx of free siderophores is perfectly balanced by the influx of free iron. Thus, beyond the boundary, *r* > *R*^*^, the concentration of free siderophores is zero. Hence, siderophore-iron complexes are only produced on the boundary *r* = *R*^*^. This defines a ‘sphere of influence’, *R*^*^ which depends on the degree of iron aggregation. For large *k*, it can reach up to 1 mm, i.e. secreting cells create a large region where no unchelated, free iron is available. The physicochemical properties of iron thus play an important role in determining the degree to which bacteria influence and modify their environment when secreting siderophores. In such regions where only siderophore-bound iron is present, other bacteria that are not able to access chelated iron could have competitive disadvantages (Fgaier & Eberl, 2010).

## C Comparing the effects of bacterial motility and siderophore secretion on iron uptake

### Increased diffusion

In the first model of contact-dependent iron uptake, the cell was assumed to be nonmotile. Alternatively to secreting siderophores, the cell could also increase its own diffusion coefficient, *D_B_*, and be motile. Potentially, if the cell moves, the chance of iron uptake upon direct encounter could be enhanced. We tested this by including a term for cell motility and measuring iron uptake in a cell relying on direct physical contact with iron sources. The direct uptake rate is approximately constant after a short initial phase and is mostly limited by the diffusion coefficient of the iron aggregates, *D_k_*, rather than the diffusion coefficient of the cell, *D_B_*,

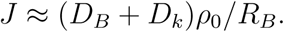

Figure S3 shows the relative increase in the diffusion coefficient the cell would need to achieve in order to equal the uptake rate of the secreter. Already for moderate aggregation levels of > 10^5^, the cell would need to increase its diffusion coefficient over 100-fold for secretion rates of 1amol/s upwards. If we use the expression for diffusion due to motility (Dusenbery, 2009),

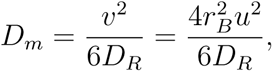

where *v* is the velocity and *u* = *v*/2*r_B_* the relative velocity of a sphere with radius *r_B_*, and *D_R_* is the rotational diffusion coefficient,

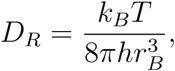

then the relative diffusion coefficient is

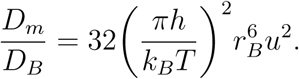

The required relative swimming speed required to gain an 100- or 500-fold increase in the diffusion coefficient would then be *u* = 2.27lengths/s and 5.07lengths/s, respectively, or a speed of around 5 − 10 μm/s.

## D Iron uptake competition

Once secreted siderophores encounter and bind iron, the chelates must successfully return to cell via diffusion. For a single cell, this process can be modelled by diffusion around a spherical sink. We generally know the probability that a chelate arrives at the cell, i.e. the hitting probability of a small diffusing particle starting at a distance *r* on a sphere with radius *R*. In one or two dimensions, *p*_1_ = 1, but in three dimensions *p*_1_ ≈ *R*/*r*. We are interested in the probability that a chelate is ‘stolen’ from a cell A by a cell B, i.e. the fraction of trajectories from the initial position O to A that pass through B. We can decompose the probability that the chelate arrives at A into direct paths, that do not pass through B, and indirect paths, that pass through B,

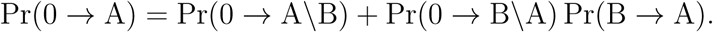

Let *p*_1_ = *R*/*r* ≈ P_r_(0 → A) be the hitting probability for a particle starting at a large distance *r* and *b* = P_r_(0 → A\B) be the probability of arriving at A by avoiding a cell B at a distance *d* (which is approximately the same as arriving at B by avoiding A for *r* ≫ *d*). Let *c* = *R*/*d* ≈ P_r_(B → A) be the hitting probability of A when starting at B (or of B when starting at A). Thus, *p*_1_ = *b*(1 + *c*), and hence *b* = *p*_1_/(1 + *c*). The fraction of chelates that would arrive at A by first passing through B is thus,

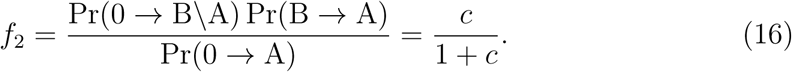

In three dimensions, for two cells at a distance *d*, *c* ≈ *R_B_*/*d*. Thus, for an immediately neighbouring cell, *d*→ *R_B_*, *c*→ 1 and *f*_2_ = 1/2, such that half of the chelates are stolen by B. However for cells that are sufficiently far apart, *d* ≫ *R_B_* and *c* ≪ 1, such that *f*_2_ ≈ *c* and the cell *de facto* no longer feels the competition of the other cell. In one or two dimensions, however, *c* = 1 for all distances and hence the competition is always 1*/*2.

## Supplementary Figures

**Figure S1:**
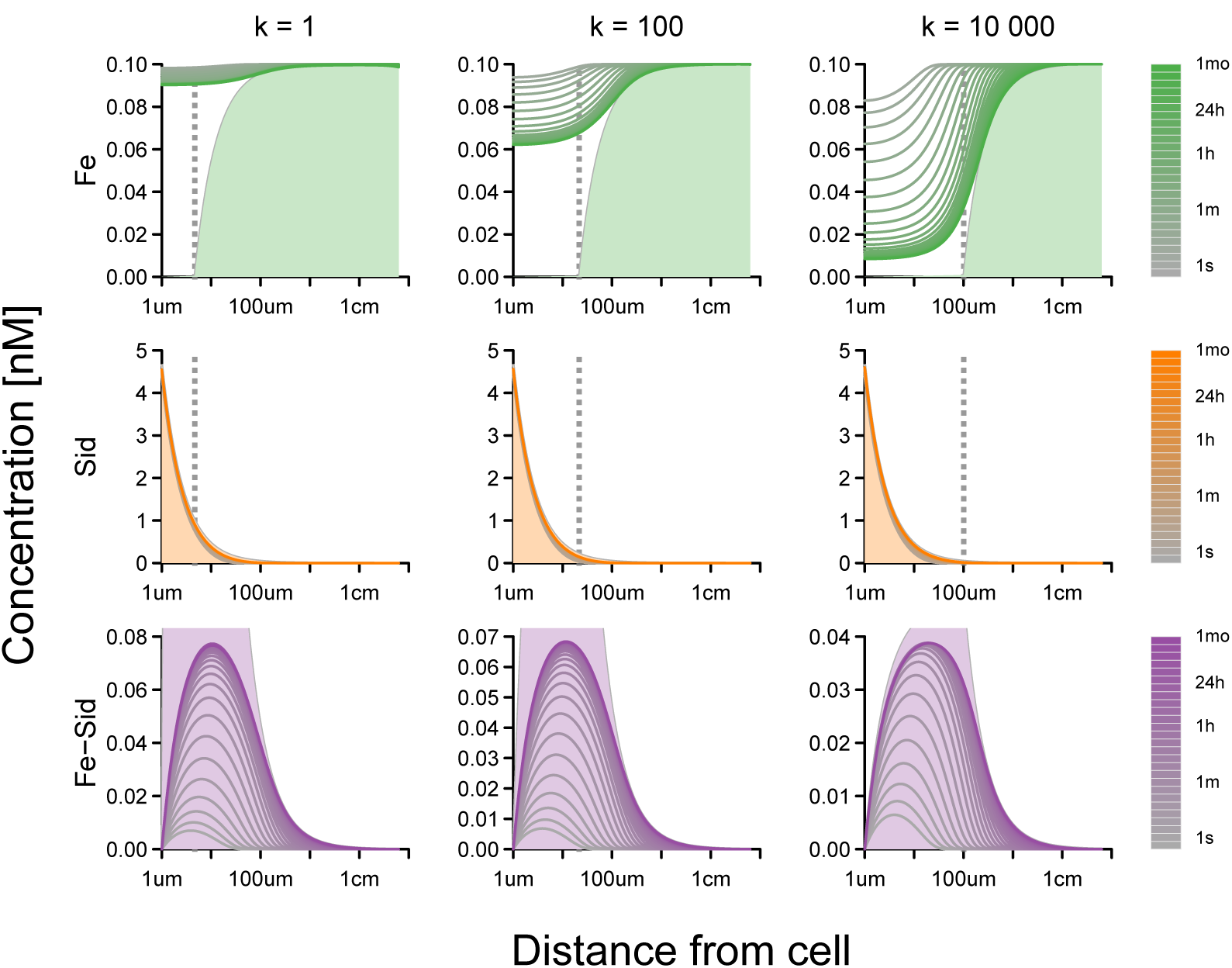
Radial distribution of free iron, free siderophores and iron-siderophore complexes over time. The figure columns correspond to different values of iron aggregation, with each aggregate containing *k* iron ions. The intensity of the color indicates the time. The vertical dashed grey lines show the large-*k* approximation of the peak iron-siderophore distribution *R\** and the shaded areas show the large-*k* approximation of the equilibrium distributions. At high *k*, the free iron close to the cell is bound by secreted siderophores and depleted. For very large *k*, the concentration at the cell surface drops to zero. The distribution of free siderophores quickly approach their equilibrium distribution, which is close to the distribution of siderophores in absence of iron. As the iron within the boundary *R*^*^ is slowly depleted, the concentration of iron-siderophore complexes first increases to high levels, before flattening out to its equilibrium distribution, where most new iron-siderophore complexes are produced at a distance *R*^*^. This equilibrium distribution, however, is only reached after a considerable time (> 1 month for *k* = 10^6^). See also Fig. S2 for *k* = 10^2^,10^6^,10^10^.

**Figure S2:**
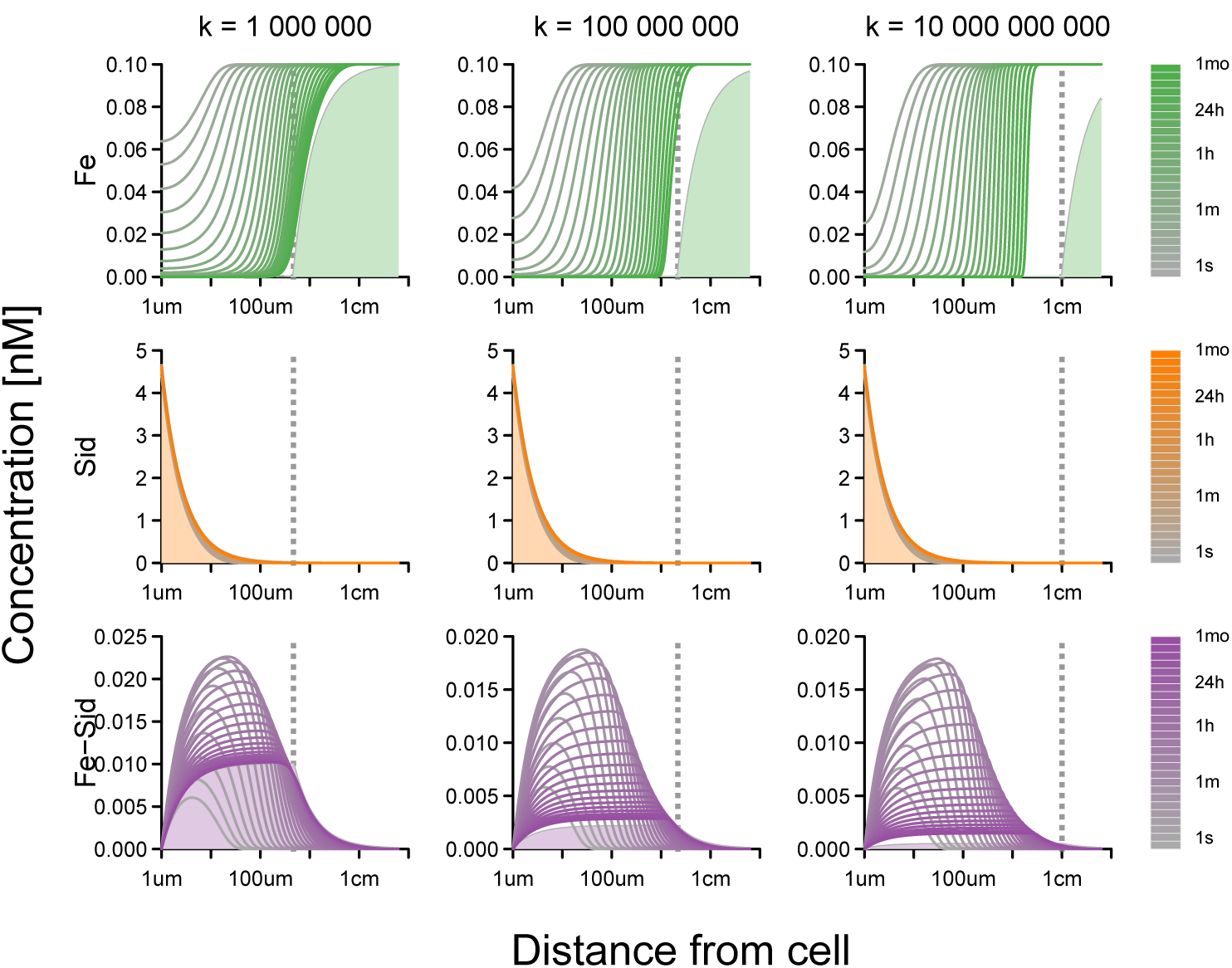
Radial distribution of free iron, free siderophores and iron-siderophore complexes over time (continued).

**Figure S3:**
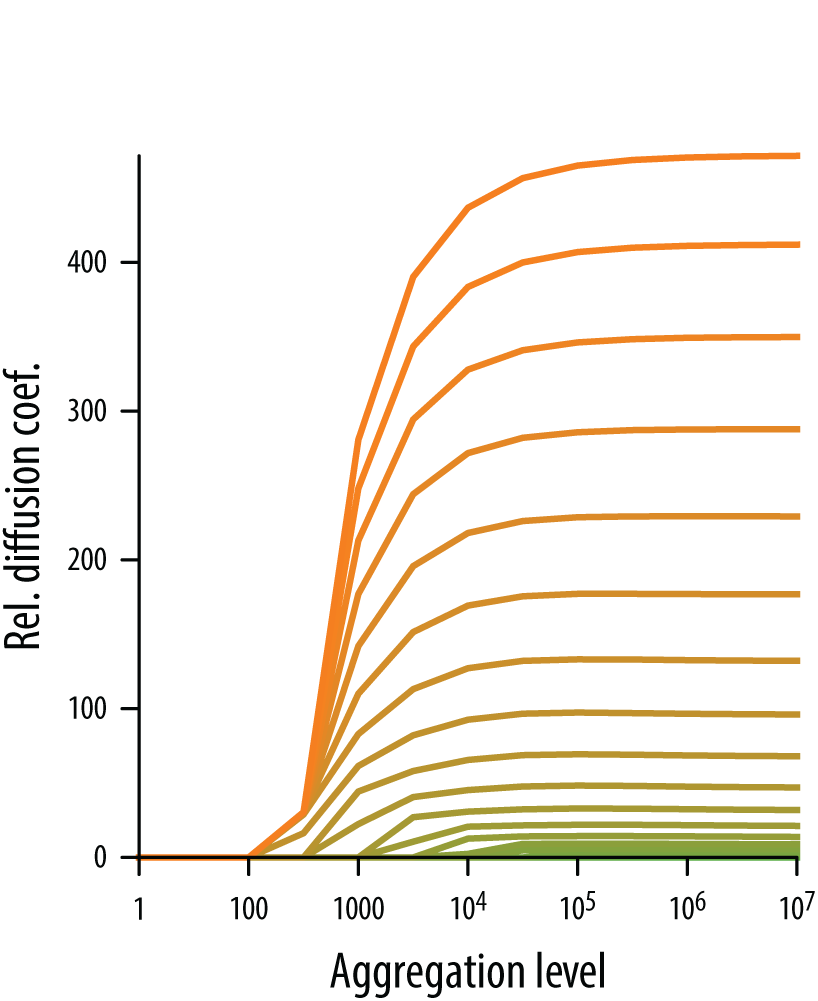
Relative diffusion coefficient required by a cell to achieve an equivalent increase in iron uptake as siderophore secretion.

**Figure S4:**
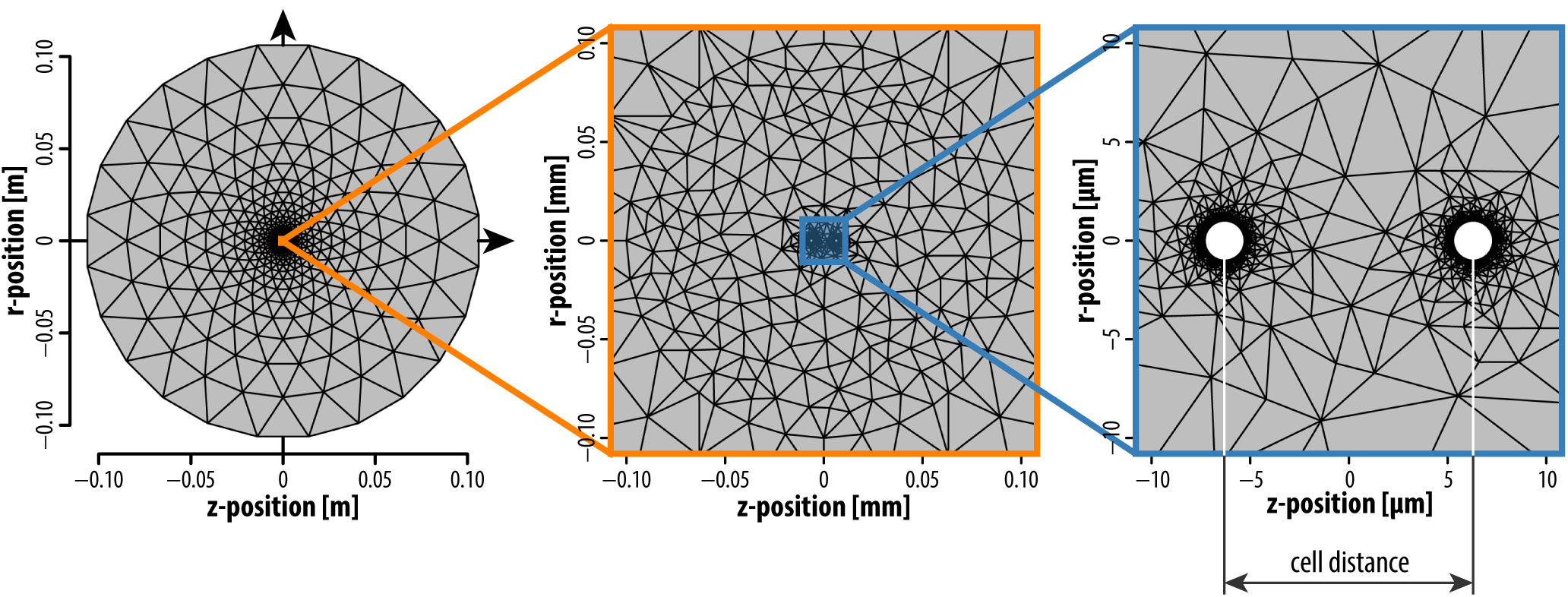
Example mesh used in the 2D numerical integration scheme. The cylindrical symmetry of two cells can be exploited to only solve the reaction-diffusion equations in two dimensions. The large length scale differences between cell spacing and siderophore diffusion require the use of an expanding mesh. A local for *r* < 0.1 mm was mesh was created and refined using the mesh generation routines in FEniCS (Alnæs *et al.,* 2015). For *r* > 0.1mm a regular circular expanding mesh was added to the local mesh.

**Figure S5:**
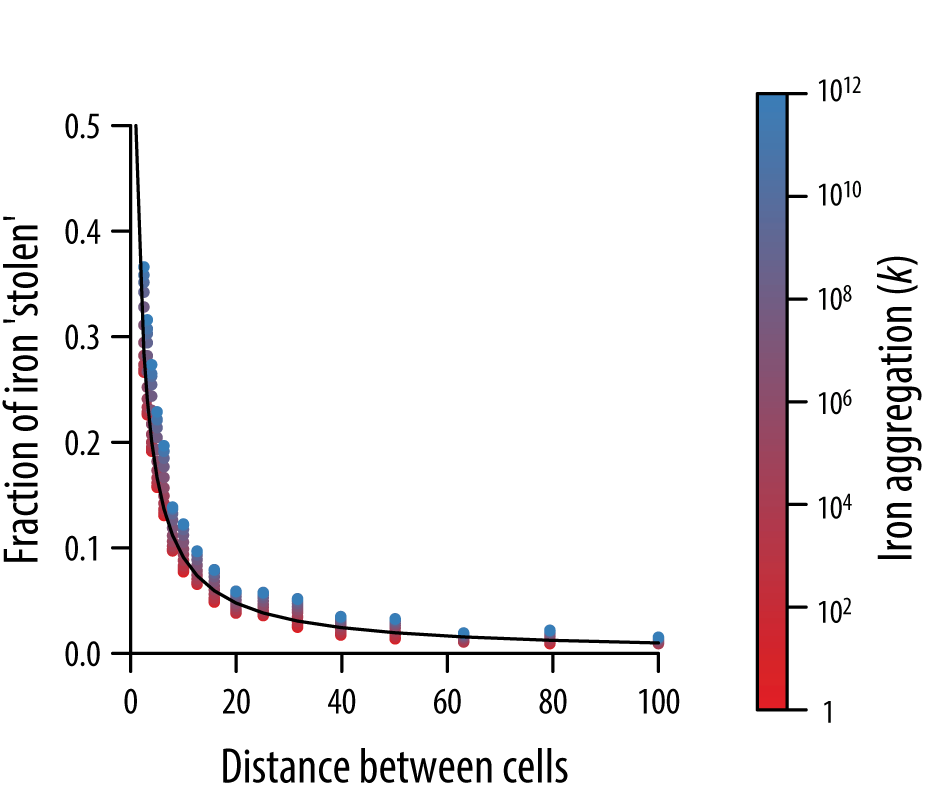
The fraction of iron that is stolen by a neighboring cell depends on the distance between cells. The approximation for the fraction of iron that is stolen by a neighbor at a distance *d* (black line) is in excellent agreement with the numerical solutions at different aggregation levels (colored points). The color scale is the same as in Fig. 3.

